# Limits to selection on standing variation in an asexual population

**DOI:** 10.1101/2023.05.11.540325

**Authors:** Nick Barton, Himani Sachdeva

## Abstract

We consider how a population responds to directional selection on standing variation, with no new variation from recombination or mutation. Initially, there are *N* individuals with trait values *z*_1_, …, *z*_*N*_ ; the fitness of individual *i* is proportional to 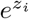. The initial values are drawn from a distribution *ψ* with variance *V*_0_; we give examples of the Laplace and Gaussian distributions. When selection is weak relative to drift 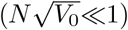, variance decreases exponentially at rate 1*/N* ; since the increase in mean in any generation equals the variance, the expected net change is just *NV*_0_, which is the same as Robertson’s (1960) prediction for a sexual population. In contrast, when selection is strong relative to drift 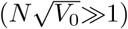, the net change can be found by approximating the establishment of alleles by a branching process in which each allele competes independently with the population mean and the fittest allele to establish is certain to fix. Then, if the probability of survival to time 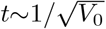 of an allele with value *z* is *P* (*z*), with mean 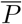, the winning allele is the fittest of 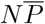 survivors drawn from a distribution 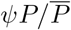. When *N* is large, there is a scaling limit which depends on a single parameter 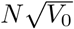; the expecte d ultimate change is 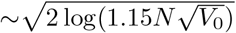 for a Gaussian distribution, and 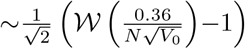 for a Laplace distribution (where 𝒲is the product log function). This approach also reveals the variability of the process, and its dynamics; we show that in the strong selection regime, the expected genetic variance decreases as ∼ *t*^−3^ at large times. We discuss how these results may be related to selection on standing variation that is spread along a linear chromosome.

---

We consider a simple and fundamental problem: how does a large asexual population respond to selection? We assume that selection acts steadily on standing variation, with no input from either mutation or recombination. Thus, the only parameters are the number, *N*, of individuals, and their fitnesses.

It is surprising that, despite its simplicity, this problem has not (to our knowledge) been addressed before. This may be because population genetics deals mainly with competition between two alleles per locus, whilst quantitative genetics has focused on genetic variance in sexual populations. Two bodies of work lie closest to the problem studied here. Robertson (1960) showed that directional selection on standing variation under the infinitesimal model causes an ultimate response, i.e., a total change in mean, equal to the genetic population size (i.e. 2*N* for *N* diploid individuals) times the change in the first generation. Moreover, he showed that the result implicitly assumes that allele frequencies evolve almost neutrally. Paixão and Barton (2016) extended this result to arbitrary epistasis, and also considered the opposite regime of strong selection. However, these results all apply to a multitude of freely recombining loci in a sexual population.

More recently, there has been a mass of work on fitness waves: the steady increase in the fitness of a population as it accumulates beneficial mutations. This work, which initially sought to explain the rate of fitness gain in serial transfer experiments with RNA viruses (Novella et al 1995), showed that the long-term dynamics of the fitness distribution in an asexual population can be described as a traveling wave with a *constant* fitness variance (Tsimring et al 1996; Rouzine et al 2003). Similar models also apply to the converse problem, of the steady accumulation of deleterious mutations through Muller’s Ratchet (Haigh 1978; Rouzine et al 2008). A key challenge is to account for the stochastic loss or establishment of very fit lineages at the leading edge of the wave, which is what ultimately maintains constant variance despite continual mutational input. Most of this work thus relies on some kind of ‘semi-deterministic’ approximation of the fitness wave, patching together a deterministic description of evolution in the bulk with a stochastic treatment of the dynamics of new mutants at the leading edge; this allows for approximate predictions for the speed of the wave, its variance and the genealogies of populations described by such waves (Desai and Fisher 2007; Good et al 2012; Fisher 2013; Melissa et al 2022). Rigorous results have been obtained only in simpler cases, typically by assuming a rather extreme form of selection (Brunet et al 2007; Berard and Gouéré 2010; Berestycki et al 2013) (though see Cortines (2016); Cortines and Mallein (2017)).

However, neither of these bodies of work apply to the transient response to selection on standing variation, in the absence of mutation or recombination. For example, consider a scenario where initially, there are *N* alleles, with some distribution of fitness. If selection were perfectly efficient, the fittest of these would fix, and its fitness would follow an extreme value distribution. However, even beneficial alleles are likely to be lost in the first few generations just by chance, and so, even with rather strong selection, the winning allele will be the survivor from this early stochastic phase. Here we show that if the initial fitness distribution is Gaussian, the winning allele is unlikely to be more than a few standard deviations above the mean, even if the population is very large; fat-tailed distributions would allow for larger but more erratic responses to selection.

Taken literally, such a model would apply to the short-term evolution of an asexual population, but it may also be used to approximate selection on alleles of a single gene, in a sexual population. When averaged over the genome of an outcrossing eukaryote, selection is likely to be weaker than recombination. However, selection may be concentrated on variation (both regulatory and coding) at a single gene. For example, in Drosophila, a 10kb region would experience recombination at a rate 10^−4^; such a region might contain hundreds of segregating sites, and hence an extremely large number of haplotypes. If the selective differences between these were greater than 10^−4^, then favourable combinations can increase intact (Sachdeva and Barton 2018). Thus, even though it neglects recombination, an asexual model may still be a useful starting point for understanding how selection acts when concentrated on small regions of a sexual genome.

In the following, we first use simulations to confirm that we can rescale time and selective value relative to the standard deviation 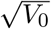 of fitness; the outcome then depends on a single parameter, 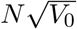 (which corresponds to *Ns* in classical models). When 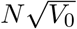 is small, vari-ance is lost mainly by random drift, over ∼*N* generations. Because the increase in mean due to selection equals the genetic variance in fitness (Fisher 1930), in this quasi-neutral parameter regime, the ultimate response is *N* times the change in the first generation, analogous to Robertson ‘s (1960) result. However, when 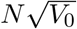 is large, variance is lost much faster, through the increase of an exceptionally fit allele that eventually fixes. We show that in this regime, the outcome can be predicted by finding the value of the fittest survivor at some intermediate time 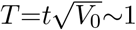, using a branching process approximation. This approach gives us both the distribution of the ultimate response, and the rate of loss of fitness variance.

## Model

The population consists of *N* haploid individuals associated with log fitness values {*z*_*i*_*}, i*= 1, …*N*, drawn from some initial distribution *ψ*(*z*) with mean zero. Individuals reproduce asexually under a Wright-Fisher model; an individual’s chance of being chosen as a parent is proportional to *e*^*z*^. For the most part, we assume that log fitness values are drawn from a Gaussian distribution with variance *V*_0_; however, we also consider heavier tailed distributions that are much more likely to generate highly fit individuals.

We neglect mutation throughout, since the aim is to understand response from standing genetic variation. Thus, in the long run, i.e., as *t*→∞, one of the alleles present at *t*=0 must fix, resulting in a net change 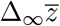 in mean log fitness, which is equal to the value of this allele. Here, we aim to understand how fixation probabilities and the dynamics of fixation depend on the initial distribution of fitness values and *N*.

### Scaling limit

This model has only two parameters, *N* and 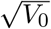, which have dimensions of time and inverse time respectively. Thus, in the scaling limit– *N* →∞, 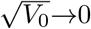, with 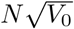 constant– evolutionary dynamics (expressed in terms of scaled time 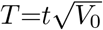 and scaled log fitness 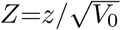) should depend only on a single non-dimensional parameter 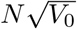. A scaling limit should exist for any initial distribution as long as it has finite variance. Moreover, in the scaling limit, the diffusion approximation should apply, and details of reproduction matter only via an effective population size; therefore, we expect that our results would extend more generally beyond Wright-Fisher reproduction.

Figure 1(a) shows the expected ultimate change 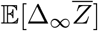 in mean log fitness as a function of 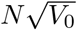 for various *N* (different symbols), where 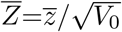 is scaled relative to the initial standard deviation of fitness. Results are shown for two different initial distributions— Gaussian (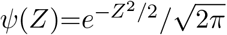; shown in shades of blue) and Laplace or bi-exponential (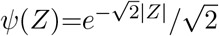; shades of red) both of which are symmetric about *Z*=0; however, our results apply more generally also to asymmetric distributions. The expectation is obtained by averaging over 10^3^-10^4^ simulation replicates for each parameter combination. Figure 1(b) quantifies the variability of net advance across replicates by plotting the the coefficient of variation (CV)— the ratio of the standard deviation to the expectation of the ultimate change— for the same parameters as in fig. 1(a). Solid black lines and dashed colored lines show analytical predictions under strong selection 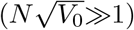 in the scaling (*N* →∞) limit and for finite *N* respectively; dotted black lines show the weak selection 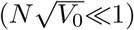 expectations (see below). Both the expected ultimate change and its CV become independent of *N* for large *N* and follow the scaling closely. However, the convergence to the scaling limit (with increasing *N*) is slower for larger values of 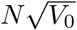 and if the initial distribution has heavier tails.

**Figure 1:**
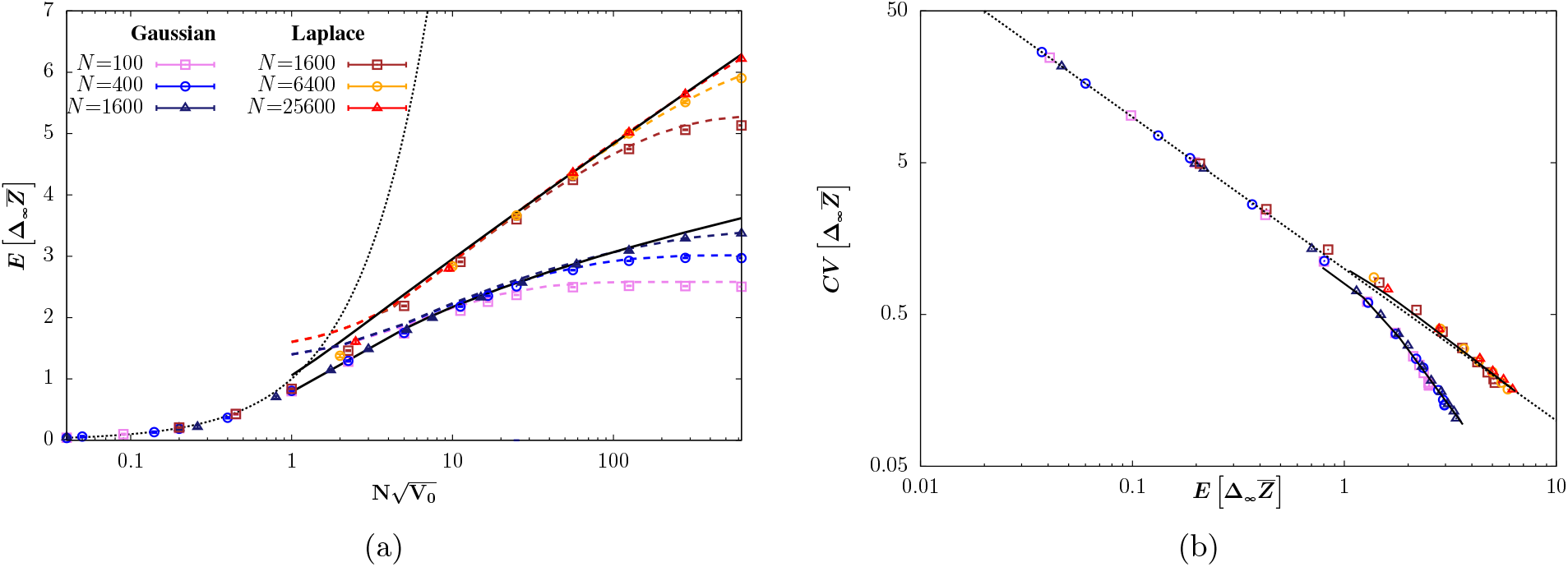
(a) Expected ultimate (scaled) change 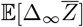 in mean log fitness and (b) its coefficient of variation (CV) vs. 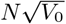, for various *N* (different symbols). Results are shown for Gaussian (shades of blue) or Laplace (shades of red) initial distributions. Symbols show simulation results; dotted black lines depict 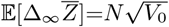 in (a) and 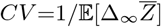 in (b), which appear to describe response under weak selection 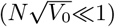. Solid black lines and colored dashed lines show the scaling (*N* →∞) and finite *N* predictions respectively in the strong selection 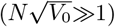 regime; finite *N* predictions for the CV are not shown since these are essentially indistinguishable from the *N* →∞ predictions (solid black lines in (b)). Scaling limit predictions for 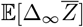 and the CV in the strong selection regime are obtained by integrating over the distribution of ultimate response in eq. (5) (and using eq. (3) to approximate the survival probability of fit alleles, with *T*_*_ set to 1 for the Gaussian and to 0.625 for the Laplace distribution); finite *N* predictions are obtained by integrating over eq. (12) in the Appendix.

Figure 1 points towards distinct behaviours of the selection response in the 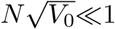 (low initial variance or weak selection) and 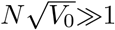 (high initial variance or strong selection) parameter regimes. We first describe these behaviours qualitatively and then outline a branching process approximation which yields surprisingly accurate predictions (solid black and dashed colored lines in fig. 1) for the distribution of the ultimate change when 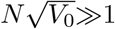.

### Response under weak selection

For 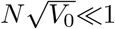, the expected ultimate (scaled) change is close to 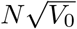 (shown by the dotted black curve in fig. 1(a)), regardless of the shape of the initial distribution. This is precisely the net advance under directional selection in a sexual population under the infinitesimal model (Robertson 1960). It can be understood in the same way, by noting that under weak selection, fitness variance decays neutrally, i.e., by 1*/N* per generation on average (see also fig. 4(b)), and that the expected change in 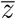 per generation is equal to the variance of *z*. Therefore, the total change in mean is 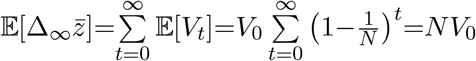; scaling relative to the initial standard deviation gives 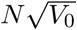.

However, unlike in sexual populations with very many loci, here (i.e., in the asexual case), the ultimate change is determined by the fitness of the single allele that fixes; it is thus highly variable across replicates, being *NV*_0_ only *on average* (for the unscaled mean). In fact, the variance of the ultimate change is approximately equal to the initial variance *V*_0_ of log fitness values, such that the variance within populations is converted entirely to variance across replicates over time. Consequently, the CV of the ultimate change is 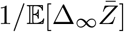 (fig. 1(b)), where 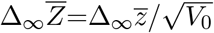 is the scaled response.

### under strong selection

By contrast, for 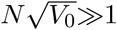, the ultimate change is much lower than 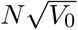 and depends strongly on the initial distribution *ψ*(*z*), being larger for longer-tailed distributions. As we show below, in this regime, selection response is governed by exceptionally fit alleles in the tail of the distribution; the fittest of these to escape initial stochastic loss is the one to fix. For instance, with a Gaussian initial distribution and for 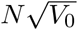 in the range 10 to 1000, the allele that eventually fixes has (on average) a fitness value between 2 and 3 standard deviations above the mean, while for a Laplace initial distribution, somewhat fitter alleles (between 3 and 7 standard deviations above the mean) fix (fig. 1(a)).

With increasing 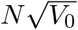, response also becomes more consistent across replicates— the CV of the ultimate change is smaller than 1 in this regime, though still non-zero in the scaling limit (fig. 1(b)). The extent of variability now depends on the shape of the initial distribution, with longer-tailed distributions resulting in higher (on average) but more variable response.

There is also a marked dependence on population size in this regime, with deviations from the scaling limit noticeable even for *N* in the tens of thousands, e.g., in the case of the Laplace initial distribution. This can be understood by noting that the ultimate change must be limited by the fitness of the fittest allele present in the population. This in turn follows a Gumbel distribution with a mean that scales as 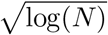 and log(*N*) for Gaussian and Laplace initial distributions respectively. Thus, for the scaling prediction to hold, log(*N*) must be sufficiently large that the fitness of the fittest allele present at *t*=0 is not limiting. A related observation (which we discuss in detail later) is that the ultimate change does nevertheless reach a scaling limit which depends only on 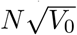, even though the fitness of the fittest allele present at *t*=0 diverges (albeit weakly) with increasing *N*.

We can understand the switch between the weak selection and strong selection regimes by noting that alleles in the tail of the distribution– with fitness values a few standard deviations above the mean– survive initial stochastic loss with a probability that is 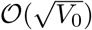 and establish, i.e., attain appreciable numbers (conditional on survival), over a timescale that is 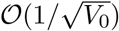. On the other hand, drift erodes diversity in the bulk of the distribution over ∼ *N* generations. Thus, the fittest surviving allele will dominate before drift has eroded diversity only if 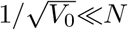, i.e., 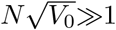; its subsequent trajectory as it outcompetes all other alleles is close to deterministic. Conversely, if the two timescales are comparable, i.e., 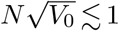, all alleles, regardless of their fitness, are affected appreciably by drift over the entire time taken for one allele to fix and the dynamics do not separate neatly into a stochastic and deterministic phase.

This is illustrated in Figure 2, which shows the distribution of fitness values at various time instants for 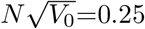 (top) and 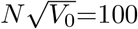 (bottom). As a reference point, the figures also show a Gaussian distribution whose variance declines over time solely due to drift (solid blue curves). In the small 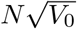 regime (top row), the fitness distribution always lies roughly within this Gaussian envelope, though there is considerable stochastic variation across replicates, especially at long times when only a few alleles survive. By contrast, in the large 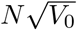 regime (bottom row), the (bulk of the) distribution is Gaussian only till *t/N* ∼ 0.01 (which corresponds here to 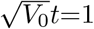), by which time a winning allele starts to emerge. In this example, the winning allele is approximately 3.3 standard deviations above the mean, which corresponds to deterministic growth by a factor of ∼*e*^3.3^≈27 during this period; however, growth conditional on survival is much faster and there are already 569 copies of this allele by *t/N* =0.01, so that it grows more or less deterministically at subsequent times and has fixed by *t/N* =0.17.

**Figure 2:**
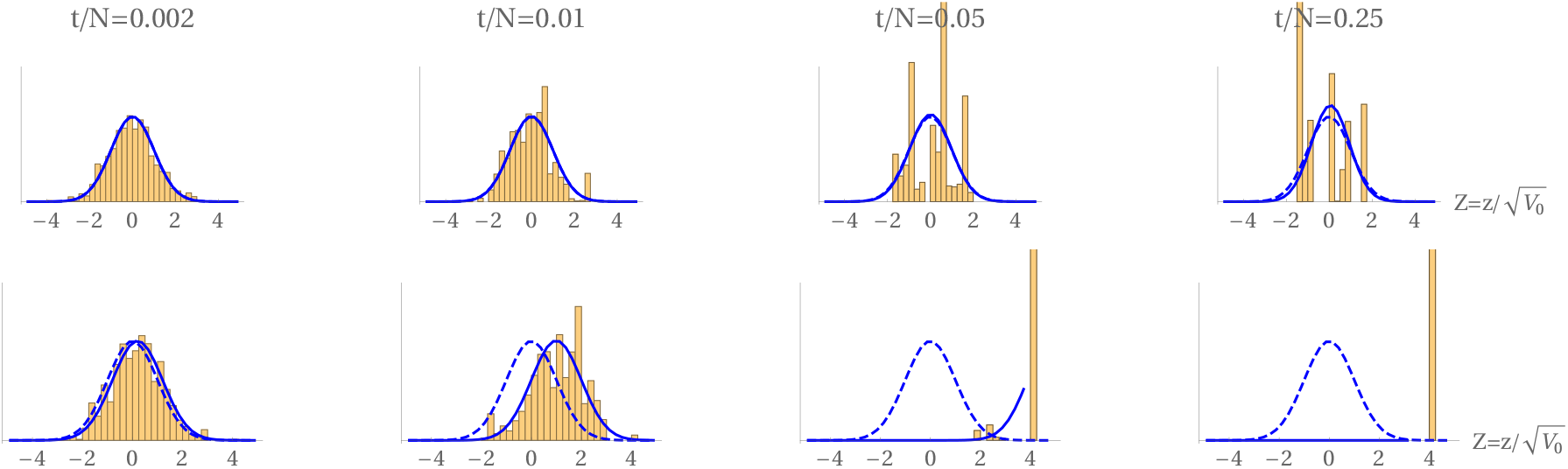
The evolution of a population of *N* =10^5^ individuals, starting from an initial Gaussian with 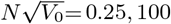, and hence variance *V*_0_=6.25*×*10^−12^ or 10^−6^ (top, bottom). The columns show the distributions of 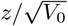 generations. The dashed curve shows the initial distribution at *t*=0; the solid smooth curve shows a Gaussian with variance that decreases due to drift, as *V*_0_*e*^−*t/N*^, and mean 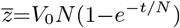.

### A branching process approximation

For 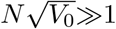, we can approximate the outcome by following the increase of each allele as an independent branching process in which an allele of value *z*_*i*_ competes with the mean 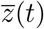 of the rest of the population. If establishment occurs over short time scales, i.e., over *t*≪*N* (as expected for 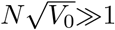; see above), then the decline in variance due to drift can be neglected, and the population mean can be approximated by 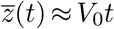. Thus, the allele experiences a time-dependent selective advantage *s*(*t*) ≈ *z*_*i*_−*V*_0_*t* during initial establishment.

Let *Q*(*t*_0_) denote the probability that an allele with advantage *s*(*t*) is lost by some time *t*_1_, conditional on it being present in *n*_0_ copies at time *t*_0_. Then, *Q*(*t*_0_)=exp[−*n*_0_*P* (*t*_0_)], where *P* (*t*_0_) is the probability of survival (until time *t*_1_) of the allele present as a single copy at *t*_0_. This form follows from the branching process approximation, which assumes that each of the *n*_0_ copies is lost independently, and also assuming *P* (*t*_0_)≪1. In the scaling limit, the change in survival probability over time can be approximated by a continuous-time diffusion process. Scaling by 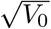, such that 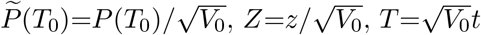 and *S*(*T*)=*Z*−*T*, the (scaled) survival probability follows the Kolmogorov backward equation:

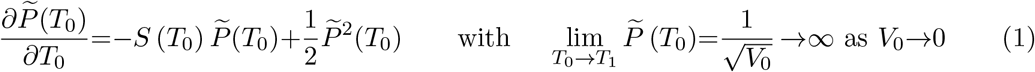

which has the solution:

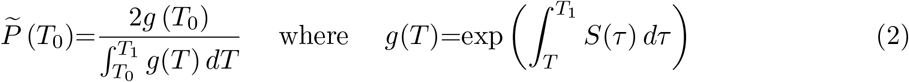

From this it follows that an allele with selective advantage *S*(*T*)=*Z*−*T*, present as a single copy at *T* =0, will survive until a time *T*_*_ with probability:

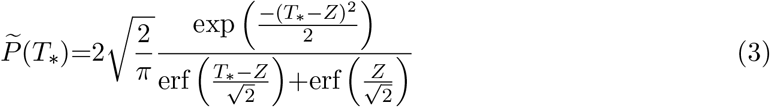

This approximation requires that the variance stay close to *V*_0_ over time *T*_*_, i.e., *t*_*_≪*N* (or equivalently, 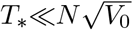) and that *T*_*_*<Z*, so that the focal allele has a positive growth rate throughout this initial phase. However, *T*_*_ must be large enough that fit alleles, having survived to this time, are very unlikely to be subsequently lost. As discussed above, both assumptions can be satisfied for some *T*_*_≈1 if 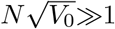.

We can obtain an even simpler approximation for the survival probability by neglecting the increase in the population mean, such that the allele has a *fixed* selective advantage *Z*. Then, the probability of survival to scaled time *T*_*_ is:

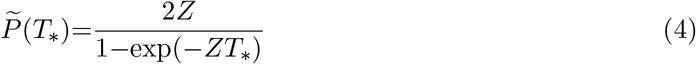

The survival probability in both eq. (3) and the simpler eq. (4) depends on the threshold *T*_*_, which is arbitrary (though 𝒪 (1), as argued above). In the following, we will explore the sensitivity of our predictions for ultimate response to the choice of *T*_*_. Surprisingly, for 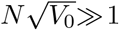, the expected ultimate change can be predicted quite accurately by using an even more drastic approximation– 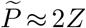, which is the asymptotic survival probability of an allele in the absence of adaptation in the bulk (see e.g., finite *N* predictions in fig. 1(a)).

### The distribution of ultimate selection response

To predict the ultimate response, we assume that the fittest allele to survive until a threshold time *T*_*_ is the one to fix; the ultimate change is then the value of this fittest survivor. First, consider a given draw {*z*} of (unscaled) allelic values; we rank these such that *z*_1_>*z*_2_>…. If we approximate the initial dynamics of very fit alleles in the tail of the distribution by independent branching processes (as described above), then each of these alleles can be associated with a survival probability *P* (*z*_1_)>*P* (*z*_2_)>…, which is given by eq. (3) (or more approximately by (4)). The probability that the fittest allele to survive up to time *T*_*_ has allelic value *z*_*i*_ or smaller, is then: 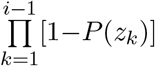, which is just the probability that all those with values larger than *z*_*i*_ are lost.

This formula gives the probability distribution of the winner from a given set of values. The fittest allele over a series of replicates will vary both because of the random initial values, and their subsequent random survival. Typically, the surviva l probability will be 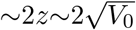, and so we expect that the winner will be ranked 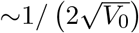. For any given draw, the best that can be achieved is bounded by the fittest allele– but the actual outcome will almost always be fixation of a much lower ranked (less fit) allele.

We now find the cumulative probability density function (CDF) *F* (*z*) of the response, by taking the expectation of 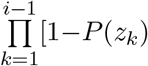 over all possible random draws of initial allelic values from a distribution *ψ*. For finite *N*, this involves first integrating over the joint distribution of {*z*_1_, *z*_2_, … *z*_*i*−1_}– the values of the top *i*−1 ranked alleles from a set of *N* iid values— over the (*i*−1) dimensional region *z* < *z*_*i*−1_ < … < *z*_2_ < *z*_1_ < ∞, and then summing over all possible *i* (see Appendix). This gives the finite *N* predictions (colored dashed lines) in fig. 1(a).

However, in the scaling limit *N* →∞, 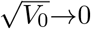 with 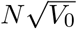 constant, there is a further simplification: the distribution of fitnesses of alleles, conditional on their having survived until time *T*_*_, is 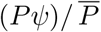 where 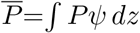 is the mean survival probability. The expected number of distinct alleles to survive the stochastic phase is thus 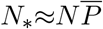, and the probability that the fittest of these has value *z* or less is:

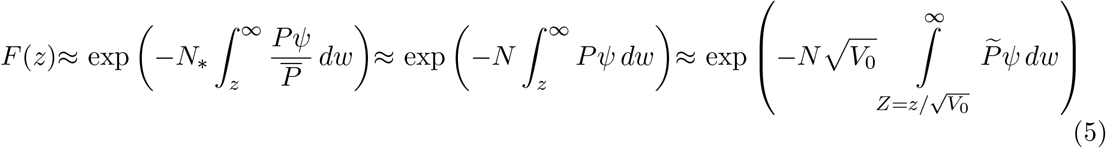

Since the probability of survival of an allele is proportional to its selection coefficient, we can write 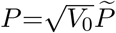 (as in the last part of the above equation). Therefore, the key parameter is 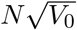, which under this approximation, only appears through the exponent in eq. (5).

The probability that the top 1, 2, …, *k* alleles to survive up till time *T*_*_ have allelic values *z*_1_, *z*_2_, … *z*_*k*_, can be recovered by differentiating ln[*F* (*z*)], and is given by [*f* (*z*_1_) … *f* (*z*_*k*_)]*F* (*z*_*k*_) where *f* (*z*)=−∂ ln[*F* (*z*)]/ ∂ *z*. Note that now *z*_1_, *z*_2_, … refer to the allelic values of the top survivors at time *T*_*_ rather than the top alleles present at *T* =0.

The theoretical predictions above depend on the unknown threshold *T*_*_— figure 3 illustrates this dependence by comparing the CDF of the ultimate change obtained from 1000 simulation replicates of a population of size *N* =40000 (red), with the predicted distribution of the fittest survivor at *T*_*_=0.5, 1, 1.5 (black, blue, purple) in the scaling (*N* →∞) limit. Solid and dashed curves show predictions that account for or neglect competition between very fit survivors and the evolving bulk; these are obtained respectively by using eqs. (3) and (4), together with eq. (5). The three sets of curves in each panel correspond to 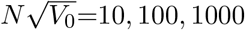 (left to right); the two panels show response for Gaussian and Laplace initial distributions.

**Figure 3:**
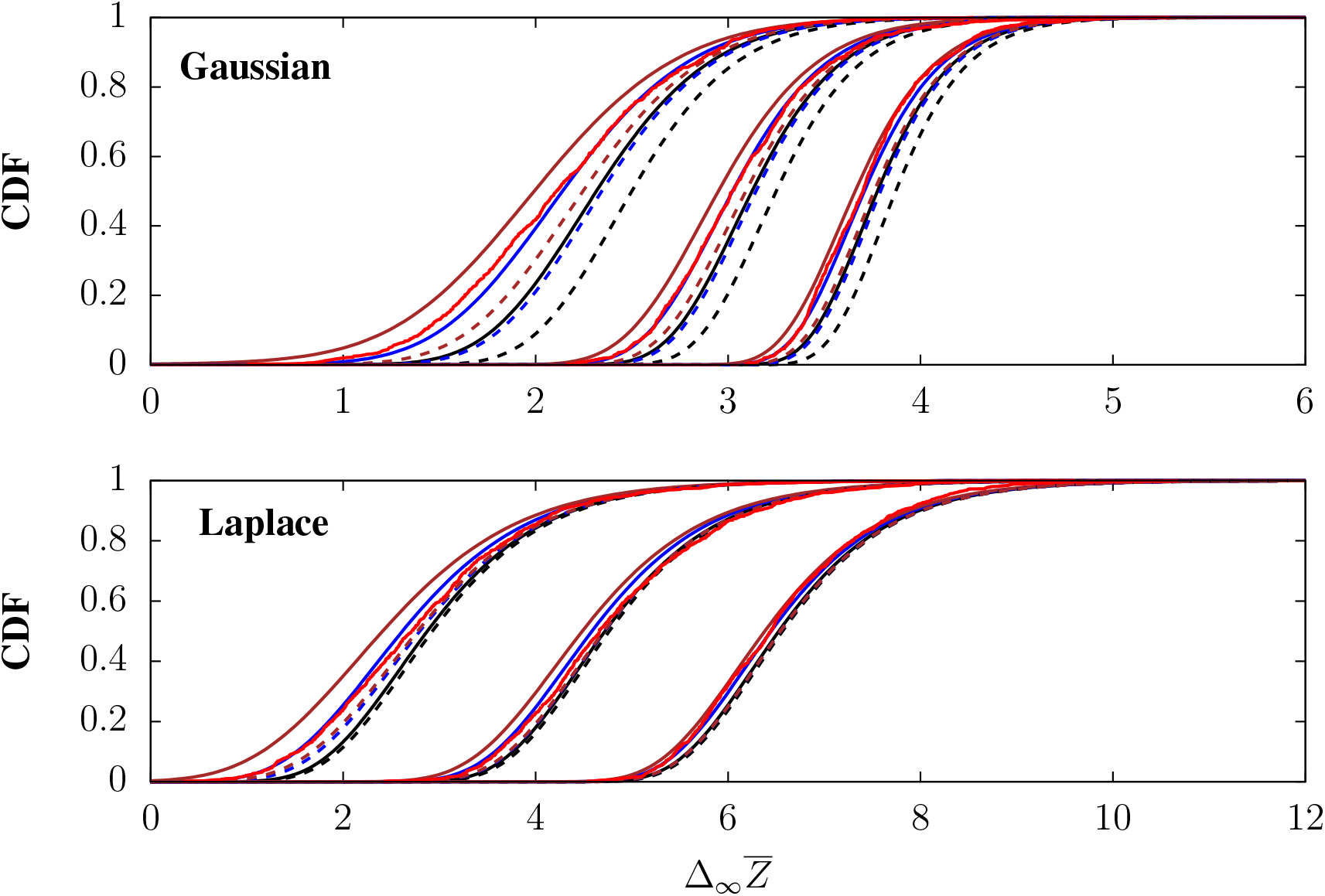
The cumulative distribution (CDF) of the ultimate response, 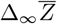, compared with various predictions. In each panel, the three sets of curves correspond to 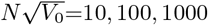; the initial distribution is either Gaussian (top) or Laplace (bottom). The CDF of 1000 replicate simulations (each initialised with a random draw of allelic values), with *N* =40000 (red) is compared with the CDF predicted by calculating the probability of survival to *T*_*_=0.5, 1, 1, 5 (black, blue, purple, solid lines right to left), from Eq. 3 (in conjunction with eq. 5). The dashed lines show the same, but using Eq. 4, which neglects the increasing mean of the bulk, and therefore predicts a somewhat higher response.

Choosing too low a value of *T*_*_ yields too high a prediction for the ultimate response since some of the alleles that have survived until a low *T*_*_ may not have established sufficient numbers and can be subsequently lost. Conversely, too high a value of *T*_*_ underestimates the aggregate selective boost enjoyed by very fit alleles during establishment (by allowing the bulk more time to catch up) and thus under-predicts ultimate response. In principle, an appropriate *T*_*_ can be estimated in a self-consistent way by assuming that it is the time at which the number of copies of the fittest surviving allele times its selective advantage crosses some threshold (such that the allele is almost certain to fix). The expected number of copies of an allele with fitness value *z*, conditional on its survival until (unscaled) time *t*_*_, is 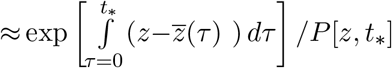 and its selective advantage at time *t*_*_ is 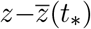, where 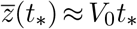 for *t*_*_≪*N*. Thus estimat-ing 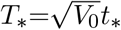 (given 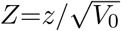) amounts to finding the *T*_*_ at which 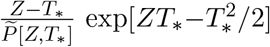 crosses some threshold. If *Z*≫*T*_*_, we have 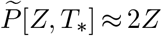, so that this criterion is approximately 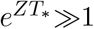. Thus, we expect *T*_*_ to scale inversely with the value *Z* of the fittest surviving allele. Approximating this value by its expectation gives 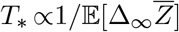, where 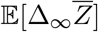 depends on 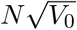 and *T*_*_.

This kind of argument can, in principle, allow us to estimate *T*_*_ self-consistently given 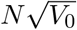. However, since 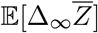 increases only weakly with 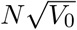 (fig. 1), in practice, predictions obtained using a fixed *T*_*_ are quite accurate across a range of 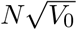 values. For example, *T*_*_∼1 appears to fit well for the Gaussian initial distribution in fig. 3, while somewhat lower *T*_*_ values (closer to 0.5) provide a better match with the Laplace distribution; this is consistent with the observation that expected response is higher by a factor of about 1.5−1.7 in the latter case. Note that the choice of *T*_*_, and more generally, accounting for adaptation in the bulk (solid vs. dashed curves) makes less difference as 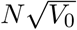 increases.

### A simple approximation for the expected ultimate response

We now find simple approximations for the expected response for both Gaussian and Laplace initial distributions. This equals 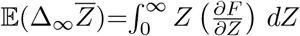 (assuming that the proportion of surviving alleles with *Z* < 0 is negligible); integrating by parts gives 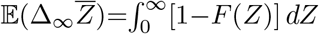. When 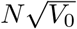 is large, the probability that the fittest surviving allele has value less than *Z* is close to zero for small *Z*, and increases sharply to 1 at some *Z* (fig. 3); thus, 1−*F* also changes sharply from 1 to 0 at this *Z*, and 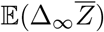 can be approximated as the value of *Z* for which 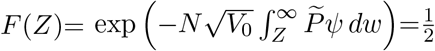, or 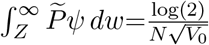. This assumes that the distribution of ultimate response is not too skewed, which appears to be a reasonable assumption, at least for Gaussian or Laplace initial distributions (fig. 3).

If we now use the crude approximation *P*(*z*)=2*z* or 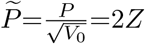, where 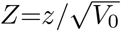, then for a Gaussian initial distribution, we have: 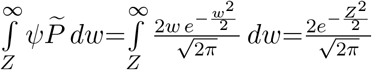. Setting this equal to 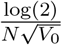 gives: 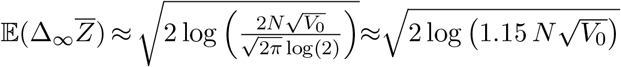. For a Laplace distribution, the same argument leads to 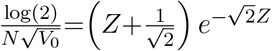; this gives 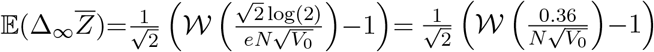,where 𝒲 [*α*] is the product log function, which is the solution to *α*= 𝒲*e*^−𝒲^. Table 1 shows that these approximations are accurate for both Gaussian and Laplace initial distributions.

**Table 1:** Mean and standard deviation of the ultimate (scaled) change as obtained from the simulations in fig. 3, compared with predictions obtained using eqs. (5) and (3) assuming *T*_*_ =1 for Gaussian and *T*_*_ =0.625 for Laplace initial distributions. The simple predictions, based on setting 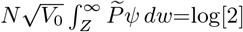, are also shown in column 4.

| Gaussian |  |  |  |  |  |  |
| --- | --- | --- | --- | --- | --- | --- |
| $N\sqrt{V_0}$ | Mean $\Delta_\infty \bar{Z}$ (sim) | Mean $\Delta_\infty \bar{Z}$ (pred) | $\sqrt{2 \log(1.15 N\sqrt{V_0})}$ | sd (sim) | sd (pred) | |
| 10 | 2.140 | 2.1729 | 2.2106 | 0.5791 | 0.5448 |  |
| 100 | 3.078 | 3.0671 | 3.0809 | 0.4222 | 0.3982 |  |
| 1000 | 3.731 | 3.7501 | 3.7546 | 0.3221 | 0.3303 |  |
| Laplace |  |  |  |  |  |  |
| $N\sqrt{V_0}$ | Mean $\Delta_\infty \bar{Z}$ (sim) | Mean $\Delta_\infty \bar{Z}$ (pred) | $\frac{1}{\sqrt{2}} \left( \mathcal{W} \left( \frac{0.36}{N\sqrt{V_0}} \right) - 1 \right)$ | sd (sim) | sd (pred) | |
| 10 | 2.84 | 2.95604 | 2.7682 | 1.1530 | 1.0547 |  |
| 100 | 4.84 | 4.83621 | 4.7102 | 1.0755 | 1.0212 |  |
| 1000 | 6.55 | 6.6603 | 6.5446 | 0.9328 | 0.9977 |  |

Thus, essentially, the mean response to selection is given by finding the expected value of the fittest of 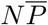 alleles, where 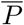 is the probability of surviving to a time 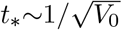 gen-erations. Surprisingly, this probability can be approximated by 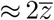, where 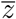 is the mean selective advantage of fitter than average alleles (i.e., alleles with *z* > 0 in case of a symmetric initial distribution) present at *t*=0, even though this neglects competition from the increasingly well-adapted bulk of the population.

### Dynamics of the fitness distribution

So far, we have focused on the initial stochastic phase which determines which allele will ultimately fix. However, the dynamics of this process are more complex and differ qualitatively in the weak vs. strong selection regimes. Figure 4 shows the variance of log fitness values as a function of scaled time, averaged over 1000 simulation replicates, for various values of 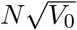 with *N* =10^4^. Time is scaled either as 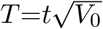 (fig. 4(a)) or *τ* =*t/N* (fig. 4(b)); the two plots are drawn on a log-log and semi-log scale respectively to illustrate the different dynamics of variance decay in the two parameter regimes. Main plots and insets show variance dynamics for Gaussian and Laplace initial distributions respectively.

**Figure 4:**
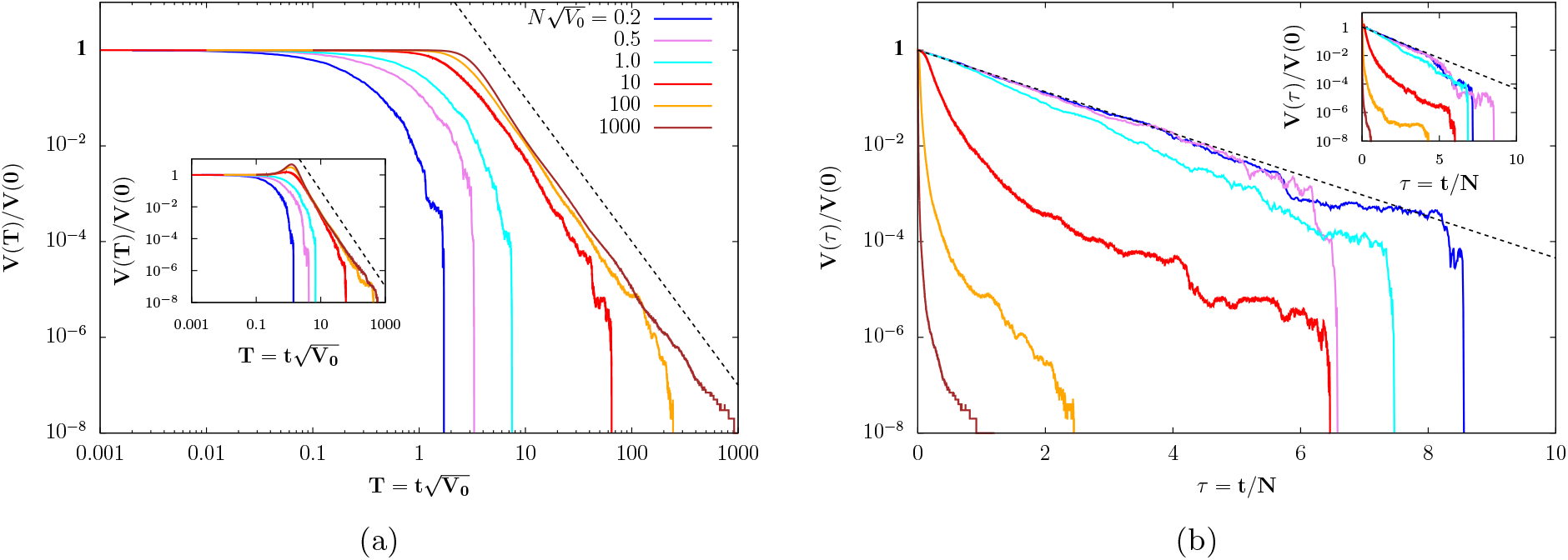
Expected variance (scaled by *V*_0_) as a function of (a) 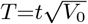 or (b) *τ* =*t/N* for various values of 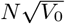 (various colors) obtained by averaging over 1000 simulation replicates with *N* =10^4^. The dashed line in (a) depicts 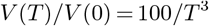 while the dashed line in (b) depicts *V* (*τ*)*/V* (0) = *e*^−*τ*^. For 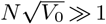, the variance decays as *T* ^−3^ independent of 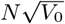, while for 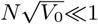, it follows the neutral expectation *e*^−*τ*^.

In the weak selection 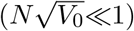 regime, the expected variance decays as *V* (*t*)≈*V*_0_*e*^−*t/N*^ (depicted via a dashed black line in fig. 4(b)), more or less independently of the value of 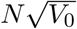 (as evident in the overlapping blue and violet curves in fig. 4(b)). This is consistent with the intuition that in this regime, the increase in frequencies of fitter than average alleles due to selection approximately balances out the decrease in frequencies of less fit alleles, so that selection has no effect on the shape of the fitness distribution (on average), and the decay in fitness variance is governed essentially by drift.

By contrast, in the strong selection 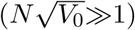 regime, the expected variance remains constant or even slightly increases (in case of the Laplace initial distribution) over a timescale 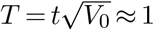, but subsequently decays as a power law ≈ *T* ^−3^ (depicted by a dashed line in fig. 4(a)). The observation of power law decay appears surprising at first glance, since in this regime, we expect long-term dynamics to be governed by the competition between the fittest few alleles to have survived the initial stochastic phase. For example, if two alleles with similar fitnesses survive, then it will take a long time for the fitter of the two to fix; assuming deterministic dynamics, variance should then decay *exponentially* at a rate proportional to the fitness difference between the two alleles.

More precisely, let *z*_1_> *z*_2_>… denote the log fitness values and *n*_1,0_, *n*_2,0_, … the number of copies of surviving alleles at some initial time *t*_*_. If dynamics are essentially deterministic for *t* > *t*_*_, then we expect alleles to grow exponentially in *relative* abundance, so that the number of copies of allele *i* at time *t* is 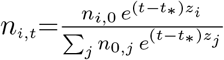. We can use this relation to predict how the fitness distribution evolves in time, neglecting stochastic fluctuations. In particular, at very long times, when only the fittest two alleles survive, we have: 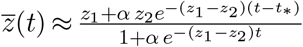 and 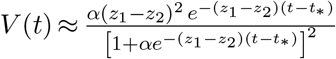, where 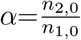 is the ratio of the numbers (at time *t*_*_) of the two fittest surviving alleles, and is itself a random variate with expectation 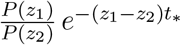, which can be further approximated by 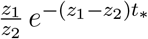 (assuming *P* (*z*) ≈ 2*z*).

As one would expect based on this simple argument, fitness variance does decay nearly exponentially in individual simulation replicates at long times (and while the less fit allele is not too rare), with a rate of decay that depends on *z*_1_−*z*_2_. The dynamics of the expected variance (averaged over replicates) can thus be predicted from the joint distribution *G*(*z, z*_1_) of the ‘gap’ *z*=*z*_1_ −*z* _2_ between the top two surviving alleles and the value *z*_1_ of the fittest survivor as: 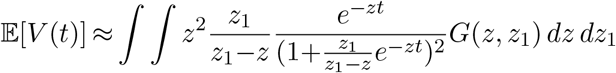. At long enough times, regardless of the exact form of *G*(*z, z*_1_) (which will depend on 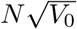 and the initial distribution *ψ*; see above), the main contribution to the expected variance comes from the *z*→0 portion of the gap distribution. In other words, as we go to larger and larger times, only replicates with smaller and smaller fitness gaps between the top two surviving alleles would still have segregating variants and only these would thus contribute to the expectation over replicates as *t*→∞. Thus, we can approximate the above expectation using Laplace’s method as:

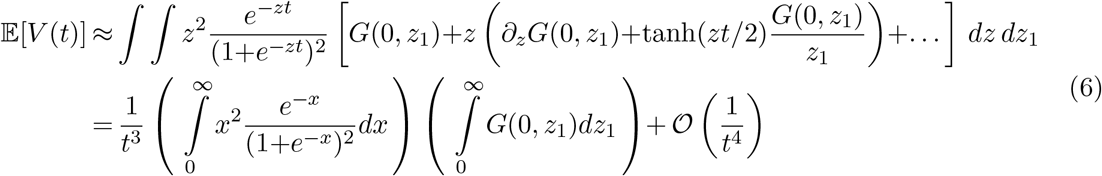

## Discussion

We consider here the limits to adaptation from standing variation in a finite asexual population, allowing for an arbitrary initial distribution of fitness values across individuals. In the absence of new mutations, the maximum possible advance under selection is limited by the fittest of *N* alleles present at *t*=0, whose fitness is (on average) ∼ log(*N*) and 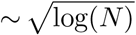 standard deviations respectively for the Laplace and Gaussian distributions considered above. However, the typical probability of survival is 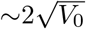, so that the actual response is governed by the fittest of 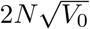surviving alleles, and is thus typically much lower than the theoretical maximum. Our derivations are an elaboration on this simple argument.

The qualitatively different— nearly neutral 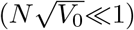 vs. strong selection 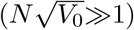 — regimes of response from standing variation in asexual populations that we identify in this study are analogous to those observed for small and large *N* in models of long-term adaptation under steady mutation, see e.g., figure 1 in Melissa et al (2022). In particular, the fact that starting with a Gaussian initial distribution, the ultimate gain in log fitness is 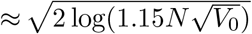 for 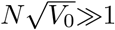 under our model, is closely related to the observation that in a population described by a fitness wave with variance *σ*, the common ancestors of future individuals are located in the leading edge of the wave, approximately 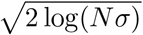 standard deviations ahead of the mean, for *Nσ*≫1 (Desai and Fisher 2007).

In this *Nσ*≫1 regime, genealogies of adapting populations are no longer described by the Kingman coalescent or its variations (e.g., with time-varying population size), but instead are strongly skewed, resulting in an excess of high-frequency derived alleles and a ‘U-shaped’ site frequency spectrum (Neher and Hallatschek 2013), as in multiple merger coalescents (Bolthausen and Sznitman 1998; Sargsyan and Wakeley 2008). One might ask how this compares with genealogies of populations adapting from standing genetic variation (as in this study) in the 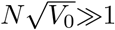 regime. In the absence of mutational input (of new alleles), genealogical structure at any timepoint during adaptation will be determined by the numbers of alleles surviving up until that point, traced back through time; these numbers, in turn, depend on the fitness gaps between surviving alleles. Thus, genealogical structure itself can change considerably during the course of adaptation; characterising these changes and the associated changes in neutral diversity along the genome remains an interesting direction for future work.

What bearing do our results have on understanding selection response from standing genetic variation in *sexual* populations? Consider an extreme scenario where variation is uniformly distributed across a linear genome, with fitness variance per unit map length *v*_*g*_. Under this infinitesimal model (with linkage), the fitness variance associated with a genomic region of map length *R* is just *v*_*g*_*R* on average. One can now ask: what is the typical map length *R*_*_ of ‘effectively asexual’ fragments that fix (under selection) without being split up by recombination in a population of effective size *N*_*e*_? If fragments evolve nearly neutrally, i.e., if 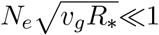, then the average time to fixation of any fragment is ≈*N*_*e*_, implying that the typical map length *R*_*_ of fragments that fix without recombining is *R*_*_∼1*/N*_*e*_. Thus, a self -consistent criterion for effectively asexual blocks to evolve nearly neutrally is that 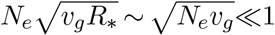.

One can also consider the opposite ‘strong selection’ regime— where fitness variance per unit map length *v*_*g*_ is sufficiently high that effectively asexual fragments can, in principle, generate sweep-like signatures as they increase under selection (Sachdeva and Barton 2018). As before, the fitness variance associated with a genomic region of map length *R*_*_ is *v*_*g*_*R*_*_; thus, under strong selection, i.e., if 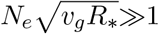, the time scale for the establishment of a highly fit haplotype spanning a map length *R*_*_ is approximately 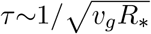(in analogy with the asexual model, where the fittest surviving allele establishes over a timescale 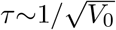). This argument assumes that fragments of map length *R*_*_ are effectively asexual over a timescale *τ*, which requires *R*_*_*τ* ∼1 or *R*_*_∼*v*_*g*_. Thus, this gives a complementary criterion— 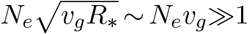 for the emergence of strongly selected effectively asexual blocks at short timescales, i.e., during *initial* response.

It seems unlikely, however, that long-term response in sexual populations (for *N*_*e*_*v*_*g*_≫1) can be understood by extrapolating from the asexual model. In particular, even if a few fit haplotypes of map length ∼ *v*_*g*_ increase intact early on, there may still be fine-scale recombination between these at longer timescales, generating new combinations of alleles, allowing for sustained response to selection in large populations. Such fine-scale recombination is expected to be especially efficient if the haplotypes that establish early on have very similar fitness, so that one of these does not rapidly fix. An interesting question is whether a long phase of competition between a few fit haplotypes (which can potentially generate new and even fitter combinations via recombination) might produce patterns in neutral diversity that mimic soft sweeps (Hermisson and Pennings 2005).

The above arguments all implicitly assume that a linear genome can be approximated as a collection of *independently* evolving asexual blocks, whereas in reality, blocks may interfere with one another as they fix. Such selective interference would be rather weak if the fitness variance per unit map length *v*_*g*_ is low, so that its effects on the dynamics of individual blocks may be captured by an appropriately defined effective population size (Robertson 1961; Santiago and Caballero 1998). However, it is less obvious whether this should hold for large *v*_*g*_, i.e., if strongly selected blocks are segregating at multiple genomic locations.

Such a caricature of the recombining genome as a mosaic of quasi-independent asexual blocks is nevertheless a useful starting point, providing at least a rough criterion for when we might distinguish selection response in sexual populations— either under a steady supply of new mutations (Neher et al 2013; Weissman and Hallatschek 2014) or from standing genetic variation (as discussed above)— from neutral evolution. These arguments all furnish self-consistently derived predictions for the characteristic map length of effectively asexual blocks (or alternatively, the typical scale over which there is linkage disequilibrium or LD) in the adapting population. This, however, obscures the fact that in reality, haplotypes that increase under selection may vary widely in length, giving rise to very heterogeneous signatures of selection along the genome. Such heterogeneity can be due to various reasons– first, variation is not uniformly spread across the genome and may at least be partly due to loci of major effect. Second, idiosyncratic patterns of LD, either in large populations or created by sampling a small number of individuals (e.g., in Evolve and Resequence experiments), may also result in a heterogeneous distribution of additive genetic variance across the genome. Importantly, this can generate considerable heterogeneity in short-term selection response along a linear genome even under the infinitesimal model (Castro et al 2019). Finally, even if the initial distribution of variance is perfectly uniform, the inherent stochasticity of evolutionary processes would still cause blocks of rather different lengths to increase during adaptation: with low initial variance (*Nv*_*g*_≪1), the resultant haplotype structure may be close to that under neutral evolution. A natural question then is whether heuristics based on asexual linkage blocks may be useful in characterizing the heterogeneity of response along the genome for *Nv*_*g*_≫1, i.e., under conditions where adaptation from standing variation produces detectable signatures in sequence data.

## Appendix

### Ultimate change under selection for finite *N*

Here, we calculate the distribution of the ultimate change under selection for 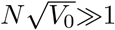 for finite *N*. As in the main paper, we assume that the fittest allele to escape initial stochastic loss and survive is the one to fix. Then, given a set of ordered and scaled allelic values *Z*_1_*>Z*_2_>… at *t*=0, and under the branching process approximation, the probability that the fittest allele to survive has rank *i* is just:

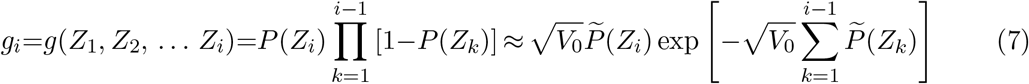

where 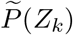 is the scaled survival probability of an allele with scaled value *Z*_*k*_; further, 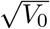 is assumed to be sufficiently small that 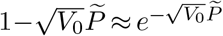. In the following, we will use the crude approximation 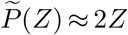 in explicit computations, but our general expression for the distribution of ultimate response (eq. (10)) does not depend on this.

The distribution of the ultimate change can now be obtained by integrating *g*_*i*_ over the joint distribution ℚ _*N*_ (*Z*_1_, *Z*_2_, … *Z*_*i*_) of the top *i* alleles present at *t*=0, and then summing over all possible values of the rank *i* of the fittest surviving allele. More precisely, the probability density ℳ (*Z*) of the (scaled) value *Z* of the fittest allele to survive the initial stochastic phase can be expressed as:

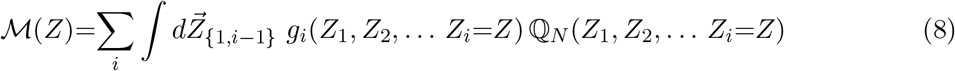

where 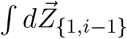 represents the (*i*−1) dimensional integral over the region *Z*<*Z*_*i*−1_<*Z*_*i*−2_*< … <Z*_2_*<Z*_1_< ∞. Note that here we work with the probability density ℳ(*Z*), rather than the cumulative density *F* (*Z*) used in the main paper.

For an initial distribution with density *ψ*(*Z*) and cumulative density function 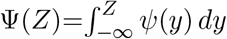, the joint distribution of the top *i* values among a sample of *N* iid values can be written as:

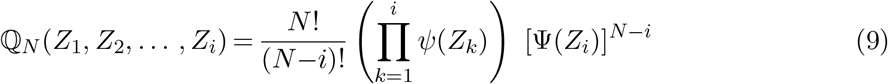

Combining equations (7),(8), and (9) yields:

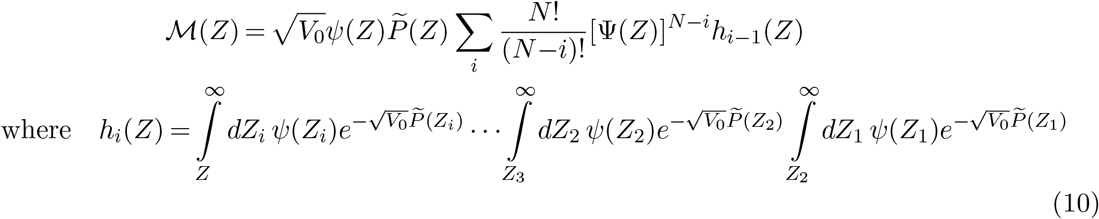

which is our main result for the distribution of ultimate change for finite *N*. The function *h*_*i*_(*Z*) is an *i*-dimensional integral; we can obtain explicit expressions for specific initial distributions by approximating the survival probability as 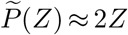 for *Z* = 0 and 0 otherwise. Then for Gaussian and Laplace initial distributions, we have:

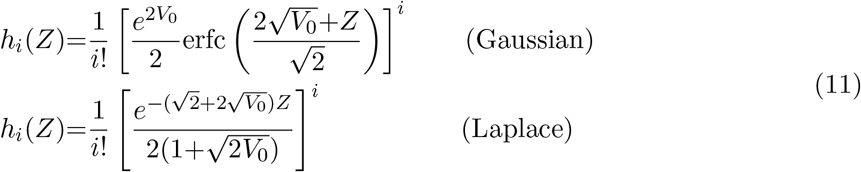

Now summing over *i*, we obtain the following approximation for (unnormalised) ℳ (*Z*):

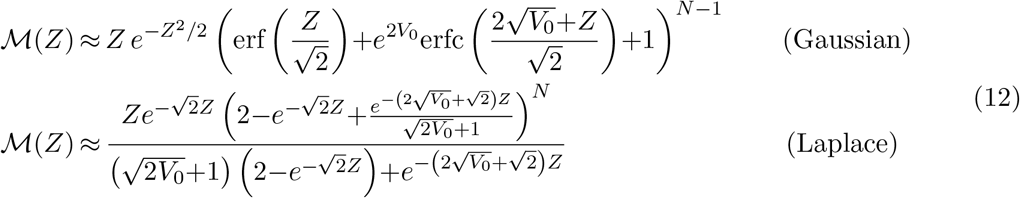

if *Z>*0, and 0 otherwise. In the limit *V*_0_→0, *N* →∞, with 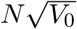 constant, the above equations simplify to the scaling limit predictions (eq. (5) in the main text).

